# Loss of Effort in Chronic Low-Back Pain Patients: Motivational Anhedonia in Chronic Pain

**DOI:** 10.1101/2025.01.09.632293

**Authors:** Samuel Alldritt, Mohammad Jammoul, Meena Makary, Susanne Becker, Daniel Maeng, Brian Keane, David Zald, Paul Geha

## Abstract

The motivational and affective properties of chronic pain significantly impact patients’ lives and response to treatment but remain poorly understood. Most available phenotyping tools of chronic pain affect rely on patients’ self-report. Here we instead directly studied the willingness of chronic low-back pain (CLBP) patients to expend effort to win monetary rewards available for wins at different probabilities and different levels of difficulties in comparison to matched pain free controls and obtained functional brain imaging on a sub-group of our sample to link behavior to brain properties. We aimed to specifically test for a differential relationship of the functional connectivity in reward and effort related brain networks, and measures of effort in patients and pain free controls. Consistent with the hypothesis of “negative hedonic shift” in chronic pain we observed that CLBP patients are significantly less willing than pain free controls to expend effort to go for high cost/high reward choices and their reported low-back pain intensity predicted increased effort discounting. Furthermore, patients’ task performance was directly correlated to functional connectivity between the ventral striatum and ventro-medial prefrontal cortex, which are major nodes in the reward processing network. Patients’ performance was not explained by their self-reported depressive symptoms. Our results present new behavioral evidence characterizing the nature of anhedonia in chronic pain and links it directly to cortico-striatal connectivity highlighting the role of this circuitry in the pathophysiology of chronic pain.

## Introduction

More than four decades ago Melzack and Casey emphasized that motivational and affective properties may be the “most important part” of the problem of pain (1). Fast forward five decades into the future, we are starting to reconceptualize the whole chronic pain experience as consisting mainly of a negative affective and motivational experience (2–6); in other words, chronic pain is not a continuum of acute pain and is characterized by distinct adaptations in the peripheral and central nervous system (7), which give rise to a negative affective condition more similar to depression, anxiety, or post-traumatic stress disorder than to an acute burn for example (8).

Chronic pain is characterized by a “negative hedonic shift” reflected in increased negative affect and decreased motivation to seek positive rewards (6). This observation is in stark contrast with the finding that acute pain administered in pain free controls increased their motivation to seek positive rewards (9). Nevertheless, the nature of the affective experience in chronic pain is still not well understood (10, 11), and the behavioral approaches to measure chronic pain affect are still lacking. Questionnaires have been developed to assess the depressive and anxiety symptoms or other negative emotional states which co-occur with chronic pain (12–14). These tools while useful in phenotyping patients (15), are mainly based on clinical observations and loosely linked to the neurobiology of chronic pain (1). In addition, while it is important to assess co-morbid depression and anxiety, these symptoms are not necessarily equivalent to chronic pain affect. Importantly, most if not all these tools were in fact developed prior to the past two decades where the role of supraspinal neural circuitries in chronic pain came into scrutiny (2, 5, 16).

Pre-clinical and brain imaging studies of chronic pain have now demonstrated the role of the cortico-striatal circuitry in tracking clinical pain intensity and affect (17–25) and in predicting the transition from sub-acute to chronic pain (26–28). Mesolimbic dopaminergic cells of the ventral-tegmental area projecting to the nucleus accumbens shell and core show region specific alterations in their firing patterns in animal models of chronic pain (29–32). The cortico-striatal circuitry mediates reward processing and is implicated in both the subjective “liking” of rewards, and the willingness or motivation to seek rewards (reward “wanting”) (33–36). Reward processing has been hypothesized to be altered in chronic pain patients since Hippocrates (37), but studies directly addressing this hypothesis in humans remain limited. We and others have demonstrated that chronic pain is associated with subjective anhedonia (4, 38–40); in addition, we and others have observed disrupted decision making when chronic pain patients are offered choices of rewarding stimuli (38, 39, 41, 42). Here we specifically test the willingness of chronic pain patients to expend effort to obtain a monetary reward occurring at different probabilities using the Effort Expenditure for Rewards Task (EEfRT) (43), which was developed to objectively measure motivation in humans. We also link distinct functional cortico-striatal circuitry involved in reward and effort processing respectively to the differences in effort expansion between groups.

## 2. Materials and Methods

### Ethics Statement

The study was approved by the Yale University and University of Rochester Institutional Review Boards and written informed consent was obtained from all participants.

### Data Sources and Participants

Data used in this work was collected at two different institutions: (1) Yale University and (2) University of Rochester Medical Center in Dr. P.G.’s lab. Between January 2018 and March 2024. Eighty-four patients with chronic low-back pain (CLBP) were recruited into the study if they had low-back pain below the 10^th^ thoracic vertebra, for at least one year with a pain intensity ≥ 30/100 on a visual analogue scale (VAS) and no other chronic pain, neurologic, or psychiatric conditions.

Therefore, patients were excluded if they reported current history of more than moderate depression, defined as a score > 19 on the Beck Depression Index (44), or history of traumatic brain injury, chronic psychiatric conditions, chronic inflammatory conditions (e.g., rheumatoid arthritis), or current ongoing chronic pain other than low-back pain. The same eligibility criteria were used to recruit 44 pain free controls who, in addition, denied any history of clinical pain. Participants completed the tasks between 9 and 11 am in the lab; after obtaining written consent a urine drug screen was obtained. Height and weight were directly measured in the lab next using a Detecto (Inc.) scale.

### Demographic and clinical data

All participants filled questionnaires to evaluate handedness, depression (Beck’s Depression Index), and anxiety (Beck’s Anxiety Index). Patients filled in addition the short-form of the McGill Pain Questionnaire (sf-MPQ)(45), and the Pain Catastrophizing Scale (PCS)(14).

### The effort expenditure for rewards task (EEfRT)

EEfRT has been thoroughly described by Treadway et al.(43). Briefly, EEfRT is a multi-trial task where participants are given an opportunity on each trial to choose between two different task difficulty levels associated with varying levels of monetary reward. Effort expenditure on this task is inversely related to anhedonia (43) and depressed patients are less willing to expend effort than pain free controls on this task (46). Each trial presents the participant with a choice between, a ‘hard task’ ((high cost/high reward (HC/HR) and an ‘easy task’ (low cost/low reward (LC/LR)) option, which require different amounts of speeded manual button pressing. For easy-task choices, subjects are eligible to win the same amount, $1.00, on each trial if they successfully complete the task. For hard-task choices, subjects are eligible to win higher amounts that vary randomly from trial to trial within a range of $1.24 – $4.30 (“reward magnitude”). The win during any task is not however guaranteed but subjects are given accurate probability cues at the beginning of each trial with high (88%), medium (50%) and low (12%) probability of win.

### Neuroimaging data acquisition sequences

Brain imaging data was available on a sub-group of subjects who performed the EEfRT and was pooled across three studies and two sites. 13 CLBP patients and 12 pain free control subjects underwent an anatomical T1-weighted scan and a resting state blood oxygen level dependent (BOLD) scan at Yale University using a Siemens 3.0 T Trio B magnet equipped with a 32 channels head-coil. The MPRAGE 3D T1-weighted acquisition sequence was as follows: TR/TE = 1,900/2.52 ms, flip angle, 9°, matrix 256 x 256 with 176-1mm slices. During the resting BOLD scan subjects were asked to stare at a cross hair; the functional acquisition sequence was as follows: TR/TE = 1,000/30.0 ms, flip angle = 60°, matrix 110 x 110 x 60 with 2 x 2 x 2 mm voxels, an acceleration factor of 4, and total number of volumes = 360.

32 CLBP patients and 18 pain free control subjects underwent an anatomical T1-weighted scan and a resting state BOLD scan at the University of Rochester Center for Advanced Imaging and Neurophysiology using a Siemens 3.0 T PRISMA magnet equipped with a 20-channels head-coil. The MPRAGE 3D T1-weighted acquisition sequence was as follows: TR/TE = 1,840/2.34 ms, flip angle, 7°, matrix 256 x 256 with 192-1mm slices. During the resting BOLD scan subjects were asked to stare at a cross hair; the functional acquisition sequence was as follows: TR/TE = 2,000/30.0 ms, flip angle = 70°, matrix 128 x 128 x 60 with 2 x 2 x 2 mm voxels, an acceleration factor of 4, GRAPPA = 2, total number of volumes = 300.

Finally, 22 CLBP patients and 6 HC subjects underwent an anatomical T1-weighted scan and a resting state BOLD scan at the University of Rochester Center for Advanced Imaging and Neurophysiology using a Siemens 3.0 T PRISMA magnet equipped with a 20-channels head-coil.

The MPRAGE 3D T1-weighted acquisition sequence was as follows: TR/TE = 1,840/2.34 ms, flip angle, 7°, matrix 256 x 256 with 192-1mm slices. During the resting BOLD scan subjects were asked to stare at a cross hair; the functional acquisition sequence was as follows: TR/TE = 1,440/32.0 ms, flip angle = 65°, matrix 104 x 104 x 72 with 2 x 2 x 2 mm voxels, an acceleration factor of 4, total number of volumes = 300.

### Neuroimaging data pre-processing

The preprocessing of each subject’s fMRI data followed the pipeline of FMRIB’s Software Library (47), www.fmrib.ox.ac.uk/fsl) and included the following: brain extraction using the brain extraction tool (BET), slice timing correction, MCFLIRT head-motion correction, and spatial smoothing using a Gaussian kernel with a full width at half maximum (FWHM) of 5 mm. Independent Component Analysis (ICA) based Automatic Removal of Motion Artifacts (ICA-AROMA) (48) was used to further remove head motion artifacts. Subsequently, additional noise reduction was performed to remove spurious variance of non-neuronal origin using linear regression. These regressors included the six parameters obtained from rigid body correction of head motion and their first derivatives and 5 components of signal from the cerebrospinal fluid following the “CompCor”-method (49). Finally, a band pass filter (0.008-0.1Hz) was applied. After preprocessing, functional scans were registered to the MNI space. Registration to high-resolution structural and/or MNI images was carried using FLIRT (50). Registration from high-resolution structural to MNI space was then further refined using FNIRT non-linear registration (47).

### Statistical Analysis

*EEfRT Data Reduction.* Because subjects could only play for 20 minutes, the number of trials completed during that time varied from subjects to subject. Time limitation of the EEfRT serves to avoid severe subject fatigue. For consistency only the first 50 trials were used. Nevertheless, there was no group difference in the number of trials between pain free controls (mean ± SEM = 67.1 ± ) and CLBP patients (70.1 ± 1.6) (p = 0.20, t-score (degrees of freedom) t (_76_) = - 1.29 unpaired t-test).

*Analysis Method 1*. The EEfRT data was analyzed following two different approaches. In the first approach, the mean proportion of the hard task choices (HC/HR) was calculated at each probability and compared across levels of probability (i.e., 12%, 50%, and 88%), or calculated at each reward magnitude and compared across reward magnitudes (i.e. < $2.5, $2.5-$3.5, and > $3.5) as within subject factors, and between groups (i.e., CLBP vs pain free controls) using generalized least squares (GLS) model with heteroskedastic error terms and autoregression 1 serial correlation (51). Age, sex, site, and years of education were included in the model as confounders.

*Analysis Method 2*. In the second approach we applied a computational model described by Cooper et al. (52) to analyze effort based decision making. Each trial of the EEfRT provides subjects with two pieces of information to consider when selecting between the high and low effort options: the reward magnitude for the high effort option and the probability of winning. To estimate the effort discounting rate (k) we fit the “full subjective value (SV) model” in which SV = RP*^h^*- *k*E, where R ($1-4.30) is the reward magnitude, P is the probability of winning, E is the amount of effort (0.3 for low effort and 1.0 for high effort), and h is the extent to which the subjects weigh subjective value based on probability(52). In this model, both *h* and *k* are free parameters. Effort perceived as extremely costly is reflected in a higher value of *k*; weighting for probability is captured by *h*. This model assumes that subjects consistently incorporate both trial-wise reward and probability when selecting. The SVs were translated into probabilities of selecting each option using the Softmax decision rule equation(53) implemented in MATLAB,

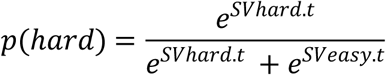

Where *t* is an inverse temperature parameter that reflects a tendency to favor options with high SVs. To test for group differences between CLBP patients and pain free controls, we compared *k* (effort discounting), *h* (subjective value), and t (inverse temperature) parameters between groups using unpaired t-test corrected for age and sex.

*ROI selection and seed-based analysis*. The seed-based connectivity analysis followed a widely applied method (54). To investigate effort-related and reward-related brain networks (55) we used the ventral striatum (vs) and the dorso-medial striatum (dms) masks as seeds, respectively, for each hemisphere of the brain. The ventral and dorso-medial striatal masks are delineated based on a functional parcellation of the striatum available from Choi et al. (56) and are illustrated in **Figure 3**. The average BOLD time course for all voxels within the regions of interest (ROIs) was extracted and corrected for brain activity in ipsilateral striatal masks outside the ROI using linear regression. This correction was performed to isolate correlations within each respective network (i.e. effort-related and reward-related) while minimizing contamination from activity in striatal regions outside the ROI. Subsequently, the correlation coefficient between this time course and the temporal variability of all brain voxels were computed using Matlab. Correlation coefficients were converted to a normal distribution using the Fisher z-transform. Because the 3 fMRI data sets originated from two different sites using 3 different MR data acquisition parameters, we applied neuroCombat o harmonize the data by removing the batch effect (57) . To study the relationship between effort expansion during EEfRT and brain connectivity we defined an anterior cingulate (ACC) mask, by using the term “anterior cingulate” in Neurosynth (www.neurosynth.org), and a vmPFC mask, by using the term “vmPFC” in Neurosynth. These masks were binarized by using a threshold z = 13 and z =10 respectively. Next, we extracted the magnitude of dms to ACC and vs to vmPFC connectivity to test if these major connections predict effort expansion by the groups. Group (i.e., CLBP and HC) average functional connectivity maps were calculated and tested against 10,000 random permutations (58), which inherently accounts for multiple comparisons, using Randomise part of FSL (59). Significant clusters were defined using threshold-free cluster enhancement (TFCE, p < 0.05) method (60), which bypasses the arbitrary threshold necessary in methods that use voxel-based thresholds.

## 3. Results

### Sample characteristics

CLBP patients had an average pain duration of 8.0 ± 1.3 years (mean ± SEM) and reported an average low-back pain intensity of 42.7 ± 2.2 on the VAS. CLBP patients and pain free control subjects did not differ in age, sex distribution, or body mass index (BMI) (**Table 1**). CLBP patients reported on average two years of education less than the pain free controls and that difference was significant (CLBP patients, 15.8± 0.3 years of education; pain free controls, 17.0 ± 0.4 years of education, t-score (degrees of freedom) t (_126_) = 2.95, unpaired t-test, p = 0.004). CLBP patients’ Beck’s Depression Index (BDI) and Beck’s Anxiety Index (BAI) scores were significantly larger than those of pain free control subjects (BDI: CLBP patients, 7.4 ± 1.0 ; pain free control, 2.1 ± 0.5, t (_93_) = - 4.0, p < 10^-3^; BAI: CLBP patients, 6.8 ± 1.1 ; pain-free controls, 2.4 ± 0.6, t (_93_) = - 3.21, p = 0.002). A sub-group of participants (27 CLBP patients and 6 pain-free controls) did not have BDI or BAI because they were part of a third study but reported mood and anxiety symptoms on the Hospital Anxiety and Depression Scale (HADS).

### Results of group comparisons across probability and reward levels in the EEfRT

We tested two GLS models corrected for age, sex, sites, and years of education for the main effect of group (CLBP vs. pain free controls), and for the interaction between group and levels of probability (model 1) and group and magnitude of reward respectively (model 2) on the preference for the high cost/high reward (HC/HR) options. Using model 1 we also found a significant effect of group on the preference of the HC/HR option (t (_376_) = 3.93; p = 0.0001), and a significant interaction (t (_376_) = - 7.08, p < 10^-4^) between group and probability levels stemming from the observation that CLBP patients expended significantly less effort than controls for options with higher probabilities of win (i.e., at 50% and 88% probability) (**Fig. 1A, S1FigA, S1Table** ). Using model 2, we found a significant effect of group on the preference of the HC/HR option (t (_376_) = - 10.73, p < 10^-4^), and a significant group by reward magnitude interaction (t (_376_) = 14.0, p < 10^-4^) (**Fig. 1B, S1Fig.B, S1Table**). Examining the graph in **Fig. 1B** we observe that CLBP patients expended less effort than pain free control subjects especially when faced with low reward magnitude but that they recovered faster as the reward magnitude increased. Because some participants had missing BMI and BDI was not collected on all participants, we repeated the GLS analysis after adding BMI and BDI in separate analyses as variables of no interest. Adding BMI does not change the results described above. Only group effects in model 1 change after adding BDI to model 1, which examines groups across the 3 levels of probability of wins; it was no longer statistically significant (p = 0.44); nevertheless, the group x probability interaction remained significant (p < 10^-4^) showing that CLBP patients were less willing to expend effort as the probability of wins increased. These analyses are presented in detail in **S2Table** with the **Supporting Information**.

**Figure 1.**
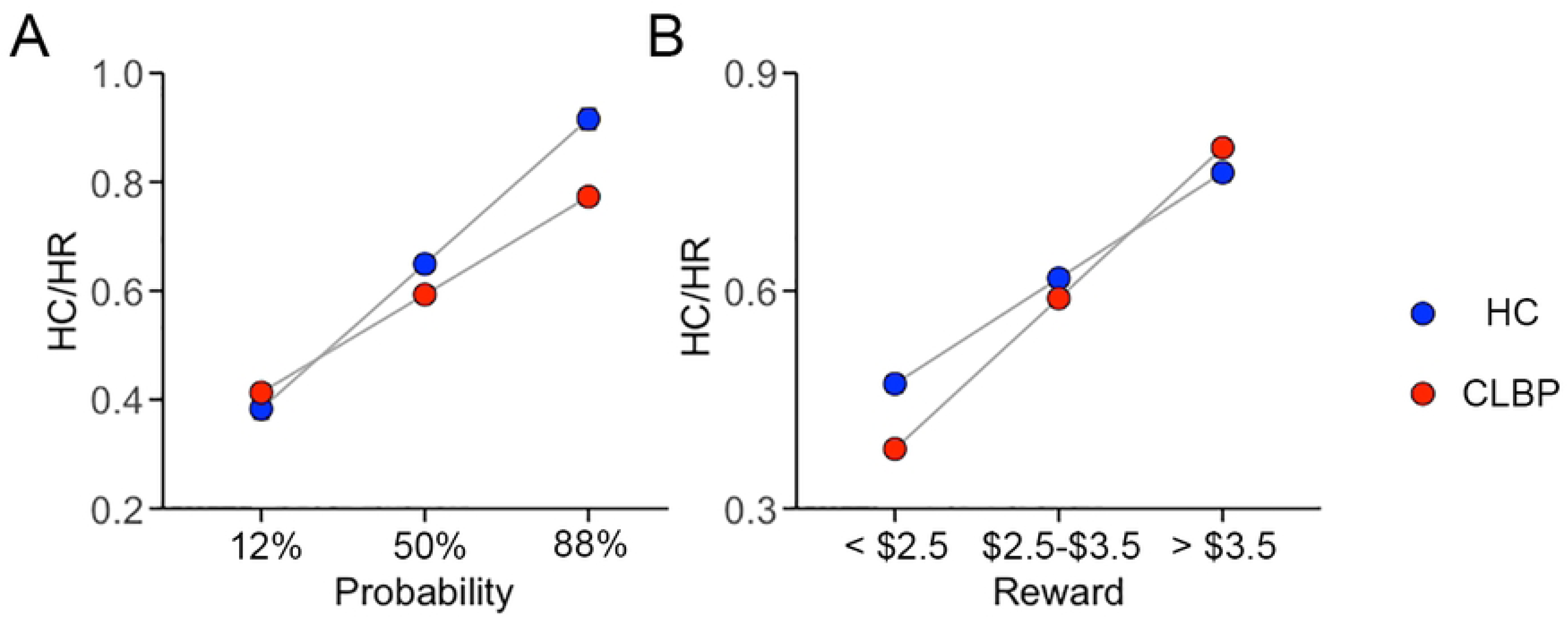
Plots showing the adjusted average ± SEM proportion of HC/HR choices (y-axis) on the first 50 trials of the EEfRT as a function of probability levels (**A**) and reward magnitudes (**B**) (x-axis) for CLBP (red) and pain free controls (blue) subjects.

### Results obtained from fitting the “full subjective value model”

The “full subjective value model”(52) was applied to obtain the parameters *k* (effort discounting), *h* (subjective value), and t (inverse temperature). None of these parameters showed significant differences between CLBP patients and pain free controls when compared after correcting for age, sex, sites, and years of education (mean *k* ± SEM in CLBP = 3.28 ± 0.41; pain free controls = 2.53 ± 0.46; p = 0.36 tested against 5000 permutations; mean *h* ± SEM in CLBP = 11.6 ± 2.3, pain free controls = 13.2 ± 4.32, p = 0.94 (non-parametric permutations); mean *t* ± SEM in CLBP = 3.4 ± 3.8 and in pain free controls = 2.4 ± 3.1, p = 0.46 (non-parametric permutations))). However, CLBP intensity reported on the visual analogue was positively correlated effort to the effort discounting parameter *k* (spearman-*ρ* = 0.24, p = 0.03) (**Fig. 2**) suggesting therefore that the more the pain the more patients were less willing to expend effort.

**Figure 2.**
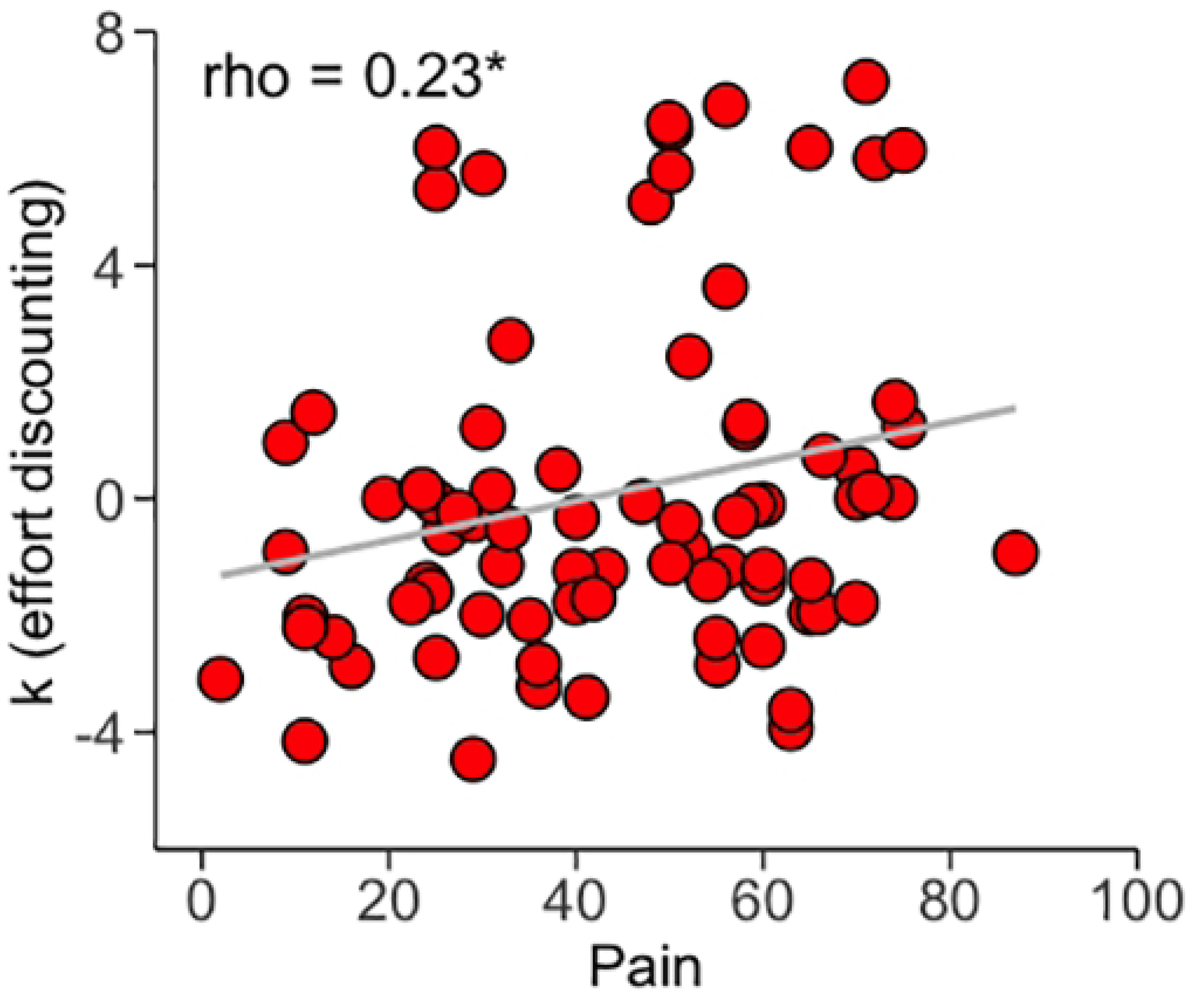
Regression plot showing how effort discounting *k* increases linearly (Spearman *rho*) with reported CLBP intensity. *, p < 0.05

### 3.4 Results of the relationship between effort and reward related brain networks and HC/HR choices

There is extensive literature on the role of ventral striatum and its connectivity to the vmPFC in encoding reward related signals (61, 62); emerging literature also highlights the distinct role of the dorsal striatum and its connectivity to the anterior cingulate cortex (ACC) in encoding effort related signals when humans seek reward (55, 63), consistent with pre-clinical findings (64). Therefore, we investigated whether connectivity within the effort-related and reward-related networks respectively as identified by (55), and calculated from BOLD images collected at rest, can predict the proportion of HC/HR chosen during the EEfRT. The effort related network was identified by using the dorso-medial striatum as a seed (**Fig. 3A**), and the reward related network using the ventral striatum as a seed (**Fig. 3B**), with both seeds derived from Choi et al. (56). Activity within each of the seed ROIs was first corrected for ipsilateral activity of the striatum remaining masks before calculating the correlation to the rest of the brain, to isolate these networks while minimizing contamination. Next, we extracted the magnitude of connection between the dms and ACC, and the vs and vmPFC where ACC and vmPFC were defined using Neurosynth.org. The right vs-vmPFC connectivity showed a highly significant positive correlation with the proportion of HC/HR choices in CLBP patients (spearman rank *rho* = 0.33, p = 0.007); the correlation in HC patients was not significant and was in the opposite direction (spearman rank *rho* = - 0.27; p = 0.11) (**Fig. 3C**). The two coefficients were significantly different between the groups (Fisher-Z = 2.82, p = 0.005). The remaining connections studied did not predict the proportion of HC/HR choices (**S3Table**)

**Figure 3.**
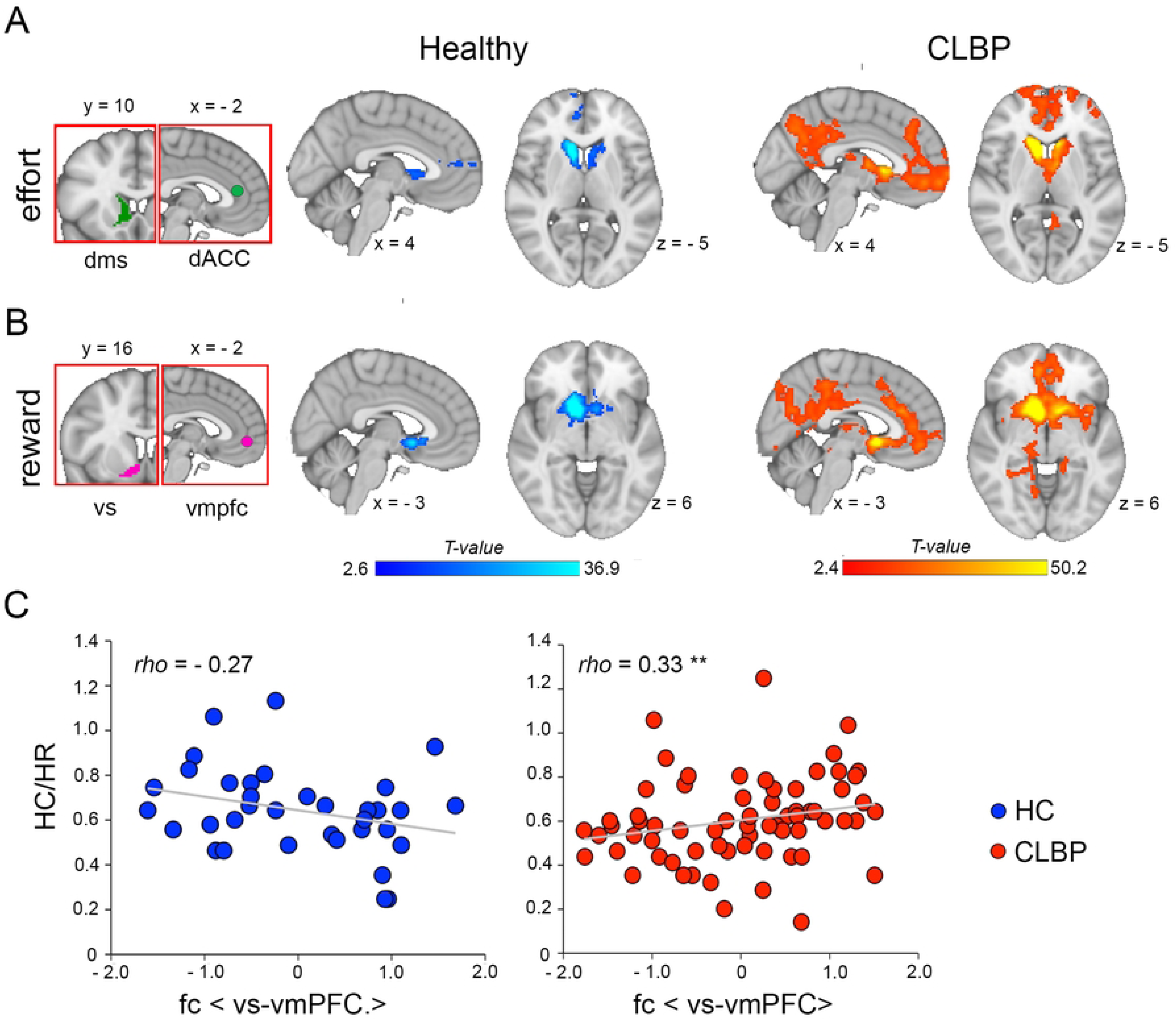
The choice of HC/HR options during the EEfRT is related to connectivity within the reward network in CLBP patients but not in HC. (**A**) Illustration of the average functional connectivity of the effort related network using the dorso-medial striatum (dms) seed in HC (blue-to-light-blue) and CLBP (red-to-yellow) subjects. The insets on the left depict the seed and the target region, the rostral anterior cingulate/paracingulate cortex (rACC/para.Cing) respectively where connectivity was queried in (**C**). (**B**) Illustration of the average functional connectivity of the reward related network using the ventral striatum (vs) as seed in HC and CLBP subjects. The insets on the left depict the seed and the target region, the ventro-medial prefrontal cortex) (vmpfc) respectively where connectivity was queried in (**C**). (**C**) Regression results showing a significant and positive linear relationship between the proportion of HC/HR chosen and functional connectivity within the reward network in CLBP patients but not in HC. **, p < 0.01.

## Discussion

In this study we present evidence that CLBP patients exhibit behavior characteristic of motivational anhedonia, as measured by an objective cost/benefit decision-making task (46). Patients suffering from CLBP are less willing to expend effort to obtain monetary rewards, even with increasing magnitude or increasing probability of wins. Furthermore, reported CLBP intensity was directly related to the effort discounting rate, supporting the observation that clinical pain hinders patients’ motivation to seek rewards. The differences in behavior between groups were also reflected by differences in how reward-related cortico-striatal brain network predicted the proportion of HC/HR choices in CLBP and HC. Specifically, the CLBP group showed stronger significant correlation than the HC group between vs and vmPFC. These observations are consistent with pre-clinical (5, 22, 24, 25, 65) and brain imaging findings showing that chronic pain patients’ motivational pathways are disrupted (2). They are also in line with our previous findings of perceptual anhedonia in CLBP patients when presented with highly palatable foods (38, 39), and reports of anhedonia (4) and disrupted emotional decision making from the literature (41, 42). This motivational anhedonia is not explained by patients’ depressive symptoms as reported on the Beck’s Depression Index. Recruitment of CLBP patients with no significant clinical depression by design may have, in fact, underestimated the loss of motivation in chronic pain, and may also explain the absence of significant differences in the parameters of the SV model (52). The selection of CLBP patients with minimal or no depression and anxiety symptoms was, however, necessary, to avoid confounds associated with the psychopathologies.

Negative affect is a major symptom of chronic pain and is often a significant negative predictor of the resolution of pain or of analgesic success (66). Nevertheless, how the affective experience of chronic pain patients differs from that of other conditions such as depressive or anxiety disorders remains unclear. Our current and previous (38, 39) results show that the loss of perceptual pleasure in experiencing or seeking rewards characterizes patients with chronic pain even in the absence of clinically significant symptoms from major affective or anxiety disorders. Consistently, Garland et al. (4), observed that anhedonia measured using Snaith-Hamilton Pleasure Scale (67) in chronic pain patients with co-morbid depression cannot be explained only by the latter diathesis. Anhedonia is conceptualized not only as a marked and consistent decrease in interest or pleasure in almost all daily activity but also loss of interest to act to seek such pleasure, and is called amotivation (68, 69). CLBP patients exhibit therefore both aspects of anhedonia independently of reported symptoms of depression and anxiety. This in turn suggests that negative affective symptoms associated with chronic pain are likely the phenotypic expression of a specific disruption of cortico-striatal circuitry distinct from the one observed in affective disorders (8). In contrast to chronic pain, acute painful stimulation delivered to pain free healthy participants increases motivation to seek monetary rewards and does not change hedonic reactions to these rewards (9). This indicates therefore that the loss of motivation observed in chronic pain patients is not simply due to an interference from a negative sensory input.

Our results, considering that EEfRT is directly modulated by dopaminergic tone (70) and that patients HC/HR choices are directly related to striatal connectivity, suggest that striatal dopaminergic transmission may also be disrupted in CLBP patients. Preclinical data clearly indicate disruptions in mesolimbic (25, 29–32) and nigro-striatal (65, 71) dopaminergic transmission in chronic pain. However, evidence in chronic pain patients is limited (72–74). A few positron emission tomography studies have reported decreased dopamine binding potential in the striatum of chronic pain patients compared to pain free controls (75), including one study on CLBP patients (72). Indirect evidence from brain imaging studies also supports this hypothesis, as CLBP patients consistently show alterations in ventral striatal activity (27). Additionally, the connectivity of this region to the vmPFC tracks back-pain intensity (17) and predicts the transition from sub-acute to chronic pain (26, 28). We have previously observed a strong and positive relationship between hedonic measures and ventral striatal (accumbens) volume in CLBP patients, but not in pain free controls, when these subjects reported their liking of highly palatable foods (39). Here, we consistently observed that motivational measures are more strongly linked to cortico-striatal (i.e., vs-vmPFC) connectivity in patients than in controls. This consistency underscores that the hedonic experience of chronic pain patients becomes closely associated with the properties of these networks as they become increasingly involved in the patients’ pain experience. This finding aligns with the established role of these circuits in cost/benefit processing (55, 76) as they integrate internal state -in this case, back pain intensity- with the cost of the effort to guide decision-making. However, a direct link between these neuroimaging findings and disrupted dopaminergic transmission has yet to be established. The observation that modulation of dopaminergic tone using dopaminergic agonists can significantly reduce pain in fibromyalgia patients more than placebo (77) and can prevent the transition to CLBP in female sub-acute low-back pain patients (78) suggest that the observed neuroimaging findings may indeed reflect a disruption in striatal dopaminergic transmission.

In addition to the dopaminergic disruption that may mediate the observed motivational deficit in CLBP patients, a disruption in opioid transmission is another potential neurochemical pathway that can explain this anhedonia. Patients with chronic pain have shown decreased binding potential of opioid ligands in the ventral striatum (79–81). Opioid receptors, which are plentiful in this sub-cortical area, (82, 83), play a significant role in pain control, hedonic processing, and negative affect as established in various studies (23, 33, 84, 85). Opioid binding in the ventral striatum is thought to contribute to hedonic encoding of rewards (86), and to the behavioral responding to reward predictive cues (87). Consistent with this role and the plasticity observed in the striatum of chronic pain patients (5, 6, 88–90) our current and previous (38, 39) work show that CLBP patients exhibit disruptions in hedonic processing whether they are asked to report their subjective ratings of pleasure or to work for rewarding stimuli like money.

Motivational and hedonic deficits are of high clinical significance because, in addition to the loss of well-being, they may lead to the emergence of other co-morbidities often observed to be associated with chronic pain such as substance use disorder (91–93), obesity (94), and depression (95). Anhedonia is associated with substance use disorders and is a target symptom for relapse prevention (3, 96, 97). Interestingly, reports of subjective anhedonia are increased only in chronic pain patients misusing opiates but not in patients taking these medications as prescribed, suggesting that anhedonia predates substance misuse (4). Anhedonia is also associated with weight gain (98, 99), and evidence suggest that this may be mechanism explaining the increased prevalence of obesity in chronic pain patients. As such, hedonic ratings predict caloric intake in hungry and satiated pain free controls, but that relationship is disrupted in chronic pain patients (38, 39).

In conclusion, our results provide behavioral evidence that chronic pain patients exhibit loss of motivation like patients diagnosed with major depressive disorder (46) . This loss of motivation cannot be solely explained by mental health symptoms. These findings contribute also to the emerging understanding of chronic pain where distinct cortico-striatal brain plasticity may be responsible for the negative affective experience observed in chronic pain patients.

## Acknowledgement

The data we collected was supported by NIDA (5K08DA037525), NINDS (R21NS1181162, R01NS127901), the Psychiatry Department at the Yale School of Medicine, and the Psychiatry Department at the University of Rochester Medical Center.

## Supporting Information

**S1 Fig.** Illustration of raw proportions of HC/HR choices for pain free healthy controls (HC) and chronic low-back pain (CLBP) patients using mean± SEM (left) and violin plots (right) while splitting the choices by probability of win (**A**) or reward magnitude (**B**).

**S1 Table**. Adjusted mean proportion of the HC/HR choices for models 1 and 2

**S2 Table.** GLS analysis of high cost/high reward choices after accounting for BMI and BDI respectively

